# The contribution of mutation to variation in temperature-dependent sprint speed in zebrafish, *Danio rerio*

**DOI:** 10.1101/2022.09.28.509995

**Authors:** Christina L. Miller, Derek Sun, Lauren H. Thornton, Katrina McGuigan

## Abstract

The contribution of new mutations to phenotypic variation, and the consequences of this variation for individual fitness, are fundamental concepts for understanding genetic variation and adaptation. Here, we investigated how mutation influenced variation in a complex trait in zebrafish, *Danio rerio*. Typical of many ecologically relevant traits in ectotherms, swimming speed in fish is temperature-dependent, with evidence of adaptive evolution of thermal performance. We chemically induced novel germline point mutations in males, and measured sprint speed in their sons at six temperatures (between 16°C and 34°C). Mutational effects on speed were strongly positively correlated among temperatures, resulting in statistical support for only a single axis of mutational variation, reflecting temperature-independent variation in speed (faster-slower mode). While these results suggest pleiotropic effects on speed across different temperatures, when mutation have consistent directional effects on each trait, spurious correlations arise via linkage, or heterogeneity in mutation number. However, mutation did not change mean speed, indicating no directional bias in mutational effects. The results contribute to emerging evidence that mutations may predominantly have synergistic cross-environment effects, in contrast to conditionally neutral or antagonistic effects which underpin thermal adaptation. However, aspects of experimental design might limit resolution of mutations with non-synergistic effects.

## Introduction

Populations can rapidly evolve in response to environmental changes (Hendry and Kinnison 1999; Hairston et al. 2005; Reznick et al. 2019), with strong evidence that local adaptation is common (Leimu and Fischer 2008; Hereford 2009). These observations point to the pervasive presence of heterogeneity among genotypes in their fitness under different environmental conditions, that is, variance due to genotype-by-environment interactions (GxE). In addition to supporting adaptive divergence of trait mean, GxE also represents the evolutionary potential of phenotypic plasticity, the expression of environment-specific phenotypes by the same genotype (Via and Lande 1985; de Jong 1990). GxE has been reported for diverse quantitative traits and environments in populations of a wide range of organisms (Des Marais et al. 2013; Wood and Brodie 2015; Saltz et al. 2018). Furthermore, environment-specific effects of segregating alleles have been characterised for specific loci (e.g., Barrett et al. 2009; Li et al. 2014). However, the interaction of evolutionary processes to generate and maintain quantitative genetic variation, including GxE, are not well understood (de Jong and Gavrilets 2000; Josephs 2018; Walsh and Lynch 2018).

Mutation is the ultimate source of genetic variation. Heterogeneous selection pressures may alter allele frequencies and shape the standing genetic variation available for on-going adaptive evolution, but the distribution of phenotypic effects of mutations that arise could ultimately determine the nature of standing genetic variation (de Jong and Gavrilets 2000; Walsh and Lynch 2018), whether populations persist (Lynch and Gabriel 1990; Gabriel et al. 1993), and how they adapt (Lorch et al. 2003; Mee and Yeaman 2019). Broadly, there are two ways that mutation can impact these processes.

First, if mutations have a consistent direction of effect on fitness across environments, heterogeneity in the magnitude of effect (strength of selection) may result in the magnitude of standing genetic variation differing among environments. In particular, for deleterious mutations, variation in the magnitude of fitness effect could result in accumulation of greater mutation load under some environments, and rapid purging under environment change (potentially leading to rapid population size declines). Martin and Lenormand (2006) and Agrawal and Whitlock (2010) reviewed studies in which mutational effects under standard rearing conditions were contrasted with effects under stressful levels of chemical toxins, nutritional resources, or temperature. No overall pattern was detected, with mutations having the same effect in both environments, or either stronger or weaker fitness effects. An updated analysis by Berger et al. (2021) arrived at the same conclusion for abiotic stressors generally, but their results suggested elevated temperatures typically increased the strength of selection.

A second way that mutational effects may impact evolutionary genetic phenomena is through their contribution to GxE associated with local adaptation. In their meta-analysis of nine studies of mutational effects, Martin and Lenormand (2006) identified a consistent pattern of increased variance in mutational effects in stressful relative to benign conditions, concluding that the frequency of beneficial mutations increased as the environmental change shifted the population further from the ancestral adaptive optimum. A shift in the distribution of fitness effects of mutations has long been hypothesised (Fisher 1930; Orr 1998), with accumulating empirical evidence of an increase in the frequency of beneficial mutations as the ancestor becomes less fit (e.g., Silander et al. 2007; Stearns and Fenster 2016). A shift in fitness effects among environments suggests mutations may typically have antagonistic pleiotropic effects (i.e., trade-off) across environments, being beneficial under some conditions, but deleterious under others, consistent with theories of local adaptation (Felsenstein 1976; Bürger 2000). In contrast, observation of predominantly positive cross-environment mutational correlations (Fry and Heinsohn 2002; Baer et al. 2006; Latimer et al. 2014) as well as direct characterisation of individual mutations under different environmental conditions (Ostrowski et al. 2005; Sane et al. 2018; Stewart et al. 2022), suggest that mutations more frequently have conditionally neutral (i.e., affecting the trait in only one environment) or concordant (synergistic) cross-environment effects.

Notably, much of the research into environmental dependency of mutational effects has focused on contrasting fitness effects between different types of environments (e.g., alternative nutrition sources) or between dichotomous, typically extreme, levels of an environmental variable. Levels of many abiotic and biotic factors vary on a continuous scale, with populations inhabiting complex, multidimensional environments. The environmental experience of any given mutation (arising spontaneously in a generation) will therefore be sampled from the distribution of natural conditions a population experiences over both its spatial range and an individual’s lifecycle. Mutations may come under fluctuating direction or strength of selection due to environment heterogeneity on these scales, which will influence the nature of standing genetic variation within populations. Studies under natural, field, conditions have revealed heterogeneity in mutational effects between temporally or spatially varying conditions (Roles et al. 2016; Rutter et al. 2018). We suggest that understanding how mutation contributes to standing genetic variation, and evolutionary potential, depends on extending our understanding of the heterogeneity of mutational effects across environmental gradients spanning ecologically relevant values.

Temperature is a key aspect of the environment, which changes over short (diurnal), medium (seasonal) and long (e.g., climate warming) time scales, as well as over small (e.g., shade vs sun patches; lake shallows vs depths) and large (latitudinal or elevational) spatial scales. Temperature affects biochemical reactions, which, in organisms with variable body temperatures (ectotherms), will affect physiological rates and all traits dependent on those rates, ultimately encompassing fitness (Huey and Kingsolver 1989; Hochachka and Somero 2002; Angilletta 2009). Temperature-dependent traits in ectotherms typically follow a stereotypical pattern of increasing values up to a so-called optimal temperature, followed by a rapid decline in values (Huey and Kingsolver 1989; Izem and Kingsolver 2005). Ecologically relevant modes of variation in this thermal performance curve shape have been identified, particularly variation in optimal temperature (hotter-colder mode) and in the width of the function (i.e., the range of temperatures over which high levels of performance are maintained: the specialist-generalist mode) (Huey and Kingsolver 1989; Izem and Kingsolver 2005). There are many examples of divergence among populations and species aligned with these modes of variation, typically reflecting known differences in thermal ecology of the taxa (e.g., Yamahira et al. 2007; Logan et al. 2018). Both hotter-colder and specialist-generalist modes of variation are consistent with antagonistic or conditionally neutral, rather than concordant cross-environment effects of contributing genetic variants (Kingsolver et al. 2001; Izem and Kingsolver 2005).

While several studies have investigated mutational effects on traits that will impact physiological processes (e.g., metabolites and enzymes: Clark et al. 1995; Harada 1995; Davies et al. 2016), few have considered whole-organism performance traits other than fitness itself (Huey et al. 2003; Ajie et al. 2005; Latimer et al. 2014). Locomotor performance is a thermally sensitive trait in ectotherms (e.g., Condon et al. 2010; Latimer et al. 2014; Logan et al. 2018), and contributes to fitness-enhancing functions such as feeding, migration, mating and predator evasion (Jayne and Bennett 1990; Irschick and Garland 2001; Husak and Fox 2008; Irschick et al. 2008; Careau and Garland 2012). While few studies have supported selection acting directly on performance indices such as maximum speed or endurance (Walker et al. 2005; Irschick et al. 2008; Wilson et al. 2020), repeated (parallel or convergent) evolution of the same performance – environment relationships (e.g., McGuigan et al. 2003; Nelson et al. 2008; Fu et al. 2013; da Silva et al. 2014; Kern and Langerhans 2019) is consistent with performance being genetically correlated with fitness. Similarly, thermal performance curves for locomotor phenotypes have diverged among taxa inhabiting different thermal environments (e.g., Logan et al. 2018) consistent with thermal heterogeneity in locomotor performance being an ecologically relevant and heritable phenotype.

Here, we applied a chemical mutagen to induce mutations in males of a laboratory strain of zebrafish (*Danio rerio*) and investigated the effects of these mutations on swimming performance of their sons. We assayed sprint swimming speed at the constant temperature experienced by these fish throughout their life (28°C; where the zebrafish stock centre system temperature is 28.5°C: Westerfield 2007) and during acute exposure to five other temperatures between 16°C and 34°C. Zebrafish are distributed throughout India and neighbouring countries, from sea level to over 1500m, and are typically found in habitats with temperatures between 16.5°C and 34.0°C, although more extreme temperatures have been reported (12.3°C - 38.4°C) (McClure et al. 2006; Spence et al. 2006; Engeszer et al. 2007; Arunachalam et al. 2013). Applying multivariate analyses and contrasting among-family variance estimated from sons of mutated sires to that estimated from sons of control, non-mutated males, we investigated the contribution of new mutation to the variation in this complex, environmentally dependent phenotype. We particularly focus on determining whether mutations have concordant effects on speed across all temperatures, or whether mutations have conditionally neutral or antagonistic effects, generating heterogeneity in mutational variance among temperatures.

## Methods

All work was conducted with approval of The University of Queensland’s Animal Welfare Unit. Adults of the Wild India Kolkata (WIK: Rauch et al. 1997) strain were imported from the Zebrafish International Resource Centre (Oregon), founding a local population at The University of Queensland, maintained for five generations prior to the current experiment (∼30 parents per sex per generation). Following protocols detailed in Rohner et al. (2011) and McGuigan and Aw (2017), males were mutagenized via a 40-minute exposure to 3mM of N-ethyl-N-nitrosourea (ENU). ENU is an alkylating agent that induces point mutations spanning a full spectrum of effects including nonsense, missense, and splice-site mutations (Knapik 2000; Wienholds et al. 2003). The ENU protocols developed for forward genetic screens to determine gene function, 1 hour exposure to 3mM ENU once a week for six weeks, induces ∼1400 – 9400 mutations (de Bruijn et al. 2009; Rohner et al. 2011). Repeated exposure to ENU saturates DNA repair mechanisms (Noveroske et al. 2000), resulting in more mutations than a single, higher concentration, dose (Hitotsumachi et al. 1985; Rohner et al. 2011). Our aim, with ∼1/9^th^ of the total standard dose, was to induce relatively few mutations, more reflective of typical long-running spontaneous mutation accumulation (MA) studies in invertebrates and plants (Halligan and Keightley 2009; Katju and Bergthorsson 2019). We consider the potential consequences of the mutation protocol further in the Discussion.

At least two weeks after ENU exposure (ensuring germline mutation transmission: Mullins et al. 1994), each mutagenized male was paired with a non-mutagenized female from the same WIK population to generate full sibling families, referred to as the Mutant treatment. The WIK stock was originally founded by a single pair of fish (Rauch et al. 1997). Reflective of this small founder size, and ongoing maintenance at a small population size (Trevarrow and Robison 2004), WIK has low polymorphism relative to wild populations (Coe et al. 2009; Suurvali et al. 2020), and long runs of homozygosity, consistent with low effective population size (Suurvali et al. 2020). Nonetheless, WIK is far from genetically homogeneous (Coe et al. 2009; Brown et al. 2012; Butler et al. 2015; Suurvali et al. 2020), and thus to infer the effect of new mutations induced by ENU against this background of standing genetic variation, a second set of males from the same WIK population, but not exposed to ENU, were also bred with non-mutagenized females, again generating full-sibling families. These Control treatment families, reared and assayed under identical conditions to the Mutant families, allowed us to estimate both the increase in heritable phenotypic variance due to new mutations, and changes in trait mean, reflecting bias in the average direction of effect of mutations.

Due to logistical constraints on rearing larvae, breeding was conducted in two blocks (3.5 weeks apart), with 27 and 23 families per treatment bred in the first and second block, respectively. Rates of natural spawning declined over time; to obtain the desired number of families (50 per treatment) and minimise age differences among families, we utilised in-vitro fertilisation (IVF) protocols on freshly collected gametes (Ransom and Zon 1999) to generate 10 mutant and four control full-sib families.

Offspring from each of the 100 families (50 per Mutant and Control treatments) were reared in two replicate 3.5L tanks, with ∼30 fish per tank at 28°C, following husbandry protocols detailed in (Conradsen et al. 2016). The 200 tanks were connected via a recirculating water system, with tank position randomised among families and mutagenesis treatment. At ∼70-100 days post fertilisation (dpf), three males from each tank (six from each family; 600 in total) were injected with a coloured elastomer tag (Northwest Marine Technology, Inc, Shaw Island, WA, USA) on either their left or right dorsal side to allow for individual identification (protocol detailed in Conradsen and McGuigan 2015). Tagged males were then transferred to new 3.5L tanks, each of which contained six Mutant and six Control males, randomly sampled from the 50 families per treatment.

### Swimming performance

Swimming speed was assayed for each of the 600 tagged males at each of six temperatures: 16°C, 20°C, 24°C, 28°C, 31°C, and 34°C. All males were first assayed at temperatures 16°C and 28°C in a semi-random order that ensured no bias in the order in which sons of the 100 families encountered the test temperatures, the time of day (morning or afternoon) they swam, or the test apparatus (two swimming flumes were used). Fish were assayed at the remaining four temperatures following the same semi-randomised design. Offspring from the first block of breeding (27 families per treatment) completed all swim trials prior to the first trails for fish from the second breeding block. In Block 1 (2), fish were 90 - 119 (114 – 149) dpf at their first trial, and 152 – 212 (183 – 219) dpf at their last trial.

Swimming speed was assayed in either a 10L (swim chamber 38cm x 10cm x 9cm length x width x height) or 30L (46cm x 14cm x 13cm) flume (Loligo Systems, Tjele, Denmark). Water in the flumes was maintained at temperatures 28°C and above using heaters (200 W or 1500 W, Quian Hu, Singapore) and at temperatures 24°C and below using chillers (440 W TECO, Ravena Italy). Fish were fasted for 12 – 24h prior to swimming. Prior to trials at non-ambient temperatures (above or below 28°C), all 12 fish from a single holding tank were placed individually in small tanks submersed in a larger tank at 28°C, which was then heated or cooled at a rate of 0.2°C per minute to bring the fish to their test temperature.

Speed was measured using a short-duration (<500 second) stepped velocity test (Brett 1964), a metric referred to as *U*_sprint_ (Starrs et al. 2011; Widrick et al. 2018). *U*_sprint_ has been shown to be strongly positively correlated with speed measured in more commonly used prolonged (>30 minute) stepped velocity tests (Reidy et al. 2000; Starrs et al. 2011), suggesting that these metrics capture similar information about individual swimming capability. The shorter-duration *U*_sprint_ was preferred in the current study due to the large number (3,600) of planned assays. To measure *U*_sprint_, fish were placed into the swimming chamber at a flow rate of 16cms^-1^ for two minutes. A pilot study confirmed this time was adequate for fish to exhibit routine behaviours. Flow was then increased by 4cm^-1^ at intervals of 15 seconds. The trial was complete (and water flow stopped) when the fish was unable to hold station and was swept to the grid at back of the chamber. *U*_sprint_ was calculated following Brett (1964) as: *U*_sprint_ = *U*_*i*_ + (*U* x (*T*_*ii*_ / *T*)), where *U*_*i*_ was the maximum velocity (cm^-1^ second) maintained for the full 15 second interval, *U* was the water speed increment (here, 4cm^-1^ second), *T*_*ii*_ was the time (seconds) the fish swam in their final velocity before tiring, and *T* was the time interval at each step (here, 15 seconds).

### Data Analyses

Two approaches have been widely used to study variation in traits that change value as a function of the environment: functional or multivariate (character-state) (Kirkpatrick and Heckman 1989; Griswold et al. 2008; Gomulkiewicz et al. 2018). Data from the current experiment, where all individuals were assayed at the same set of temperatures, is well suited to a multivariate approach (Gomulkiewicz et al. 2018). Implementing a multivariate analytical approach also directly supported comparison to results from the only other study, that we are aware of, investigating how mutational effects on locomotion vary with temperature (Latimer et al. 2014), and allowed us to explicitly address questions about heterogeneity in the magnitude of genetic (mutational) variance among temperatures, as well as the correlation in mutational effects across temperatures.

Swimming speed (*U*_sprint_) was assayed for a total of 594 males, with data at all six temperatures available for 576 of these. Estimates of quantitative genetic parameters are sensitive to extreme (outlier) values. Here, 27 observations of *U*_sprint_ were greater than 3.0 standard deviations (SD) from the mean and excluded from all analyses. If these observations reflected large effect mutation, their exclusion would under-estimate mutational effects. However, outlier individuals were widely distributed across 22 families, 12 Control and 10 Mutant, suggesting that they do not reflect mutational effects.

We first investigated whether induced mutations affected either the mean speed, or the average relationship between temperature and speed. We fit the following model using maximum likelihood within PROC Mixed in SAS (SAS Institute Inc. 2012):

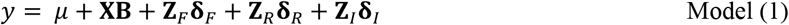

where y was the vector of *U*_sprint_ observations, *μ* was the global mean *U*_sprint_ and **X** was the design matrix relating observations to their level of the categorical fixed effects, **B**. The fixed effects of interest were mutation treatment (Mutant or Control), temperature, and the interaction between mutation treatment and temperature. Several other fixed effects were fit to account for additional potential sources of variation in speed. Trial, the six repeated measures of speed per individual, captured variation due to age, experience, and any changes in general laboratory conditions over the duration of the experiment. Time, categorised as AM or PM, was fit to account for potential diurnal effects on speed. Flume and block were fit to, respectively, account for differences in swimming performance between the two different flumes (10L versus 30L), or breeding blocks (containing fish from 27 and 23 families per treatment, respectively).**Z**_*F*_, **Z**_*R*_ and **Z**_*I*_ were the design matrices for the variance in speed attributable to the random effects of Family, replicate Rearing tank (nested within Family), and Individual (nested within tank), respectively. Reflecting the repeated measures nature of the experimental design, these random effects were each modelled as unconstrained (co)variances in speed around temperature-specific intercepts (**δ**_*F*_ **δ**_*R*_ and **δ**_*I*_)To assess the null hypotheses of no effect of mutation treatment on mean speed or temperature-specific speed, the Satterthwaite degrees of freedom correction was applied. Among-family variance was heterogenous between mutation treatments (see Results), and we investigated other models to ensure results were robust. We modified model (1) to fit treatment-specific random effects; to accommodate zero estimates of the hypothesis *F*-ratio denominator (due to no among-family variance in Control treatment), we applied a log-likelihood ratio test to the nested models in which the fixed effect of interest was fit versus not fit. The interaction between mutation treatment and temperature was tested first, and this no-interaction model was the reference model against which the main effects of mutation treatment and temperature were assessed. Both approaches supported the same conclusion and we therefore, report only the results from model (1).

Second, we investigated how mutagenesis affected the among-family (co)variance in swimming speed across the six temperatures. Mutagenesis of sires can contribute to differences between the Mutant and Control populations in both the among-family (where brothers inherit the same mutation from their father) and within-family (where brothers inherit different mutations) variance. However, variation between the two replicate rearing tanks per family, and among the three brothers sampled from each tank will reflect not only these genetic differences, but also the micro-environmental variation between and within tanks, respectively, which cannot be further partitioned out given our design. Therefore, we focus our investigation on among-family variation, which can be unambiguously assigned to genetic causes.

To estimate among-family variance, data was first centred (mean = 0) on the respective level of each of the fixed effects included in model (1). This approach is equivalent to fitting fixed effects in the analysis but improves model efficiency. Using REML in PROC MIXED to fit a modified version of model (1) (no fixed effects) to the data for each mutation treatment separately, we applied a factor analytic modelling approach (Hine and Blows 2006) to determine the statistically supported axes of the among-family covariance matrix. The among-family variation in swimming speed was constrained to zero (no among-family variance, implemented by not fitting the among-family effect) through to six dimensions (implemented using TYPE=FA0(*n*), where *n* was the number of dimensions, ranging from 1 to 6). A log-likelihood ratio test (LRT) was applied to test whether adding a dimension improved model fit; the difference in log-likelihood between nested models follows a chi-square distribution with the degrees of freedom equal to the difference in the number of estimated parameters.

We then analysed data from both treatments within the same model, estimating treatment-specific random effects (implemented using the GROUP statement), to test the null hypothesis that mutagenesis affected genetic variation. We used a LRT to compare a model in which treatment-specific among-family variance was estimated to a model in which a common, pooled, among-family variance was estimated; given evidence of low-dimensionality (see Results), this model was fit with one dimension of among-family variance, but results were consistent for higher-dimension models.

To further investigate the nature of the among-family variance in speed, we estimated the unconstrained (TYPE=UN) treatment-specific among-family covariances. We placed robust confidence intervals on model estimates using the REML-MVN sampling approach (Meyer and Houle 2013; Houle and Meyer 2015). We used the MASS package (Venables and Ripley 2002) in R (R Core Team 2020) to draw 10,000 random samples from the distribution 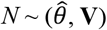 where 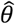 was the vector of covariance parameter estimates, and **V** was the asymptotic variance-covariance matrix from the REML model. While the REML variance estimates were constrained to be positive, the REML-MVN samples were not (i.e., were on the G-scale: Houle and Meyer 2015). We therefore interpreted the statistical significance of individual parameter estimates based on whether the confidence intervals (CI) included zero, which is equivalent to applying a LRT (Dugand et al. 2021). For variances, this is a one-tailed test (as variances cannot be negative; 90% CI), while for covariances it is a two-tailed test (95% CI). We used the ‘eigen’ function in base R (R Core Team 2020) to decompose the unconstrained REML estimates of among-family covariance to their major axes, and projected these axes (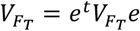 where 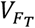 is the among-family variance in treatment T, *e* is an eigenvector of the REML among-family matrix, and ^*t*^ indicates the transpose) (McGuigan and Aguirre 2016) through the 10,000 REML-MVN samples to place CI on the eigenvalues.

## Results

Mutation did not change mean sprint speed (main effect of mutation treatment: *F*_1, 99.8_ = 0.31, *p* = 0.5767), or the response of sprint speed to temperature (mutation treatment x temperature interaction: *F*_5, 99.3_ = 0.48, *p* = 0.7909) (Figure 1A). *U*_sprint_ depended on temperature (main effect of temperature: *F*_5, 115_ = 1861.76, *p* < 0.0001), exhibiting the classical pattern of thermal performance curves, with a rapid increase in speed until ∼28°C (maintenance temperature), although there was little decline in speed by the maximum assayed temperature (34°C) (Figure 1).

**Figure 1.**
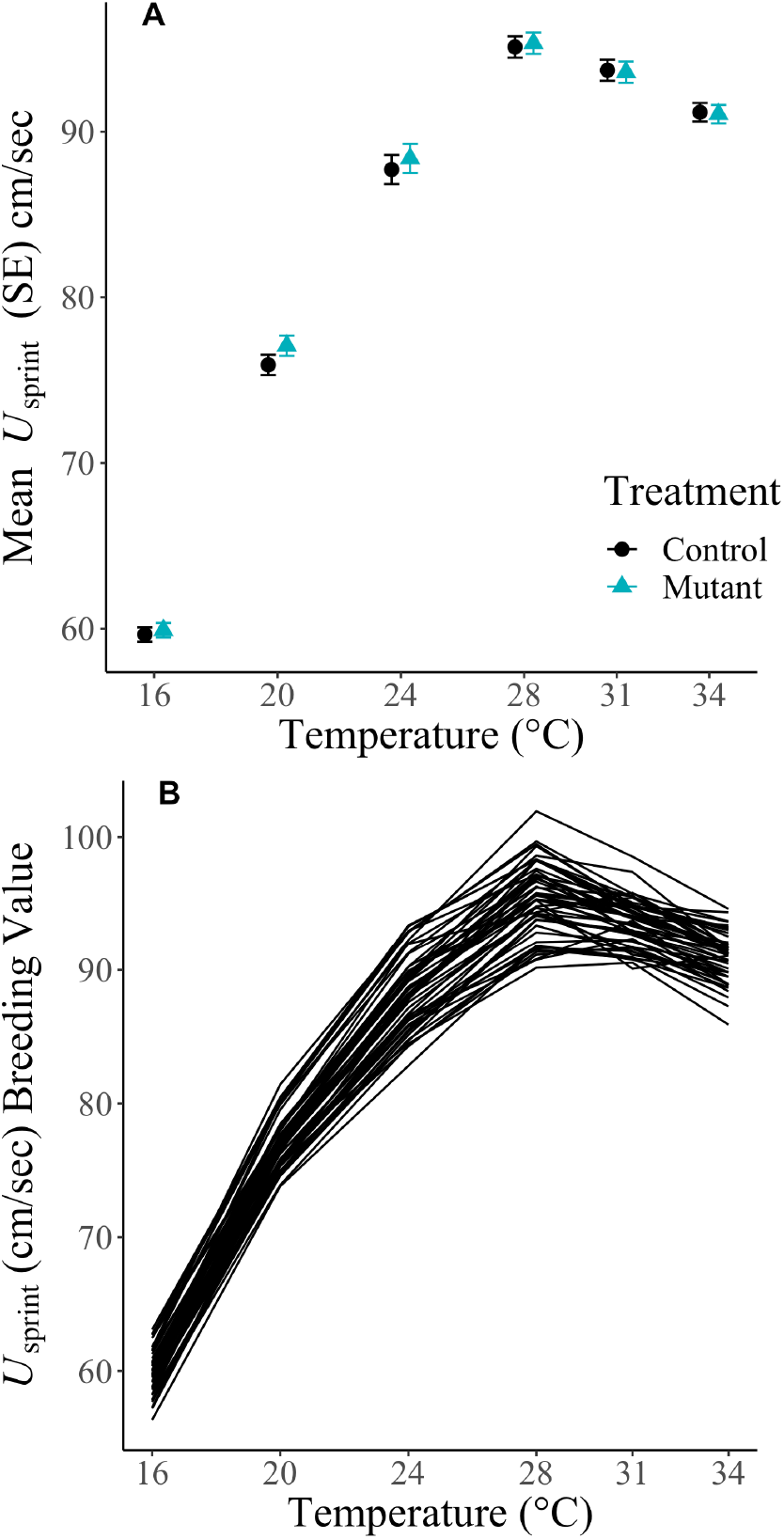
Variation in swimming speed with temperature. A) Treatment mean swimming speeds. Plotted are the least squares means (± SE) from model (1) at each of the six temperatures for the Control (black circles) and Mutant (blue triangles) treatments. B) Among Mutant family variation in swimming speed. Plotted are the Best Linear Unbiased Predictors (BLUPs) at each of the six temperatures from model (1) (fit to centred data) for the Mutant treatment, with lines connecting the point estimates for each family. The Control treatment BLUPs are not presented as they were invariant at most temperatures (Table 1).

In the Control treatment, the estimated among-family (genetic) variation in swimming speed was zero at four of the six assayed temperatures, ranging up to a maximum of 3.08 (Table 1A). Consistent with expectations, among-family variance was greater in the Mutant treatment at every temperature, ranging from 2.55 up to 11.70, although only at 24°C was the among-family variance statistically distinguishable from zero (Table 1B). We rejected the null hypothesis that Mutant and Control treatments had the same among-family variance (Log-likelihood ratio test of fit of models estimating one-dimension of treatment-specific versus pooled among-family variance: *χ*^2^ = 13.63, df = 6, *P* = 0.0341). Thus, the data supported the hypothesis that the ENU mutagenesis introduced new genetic variation in swimming speed.

**Table 1.**
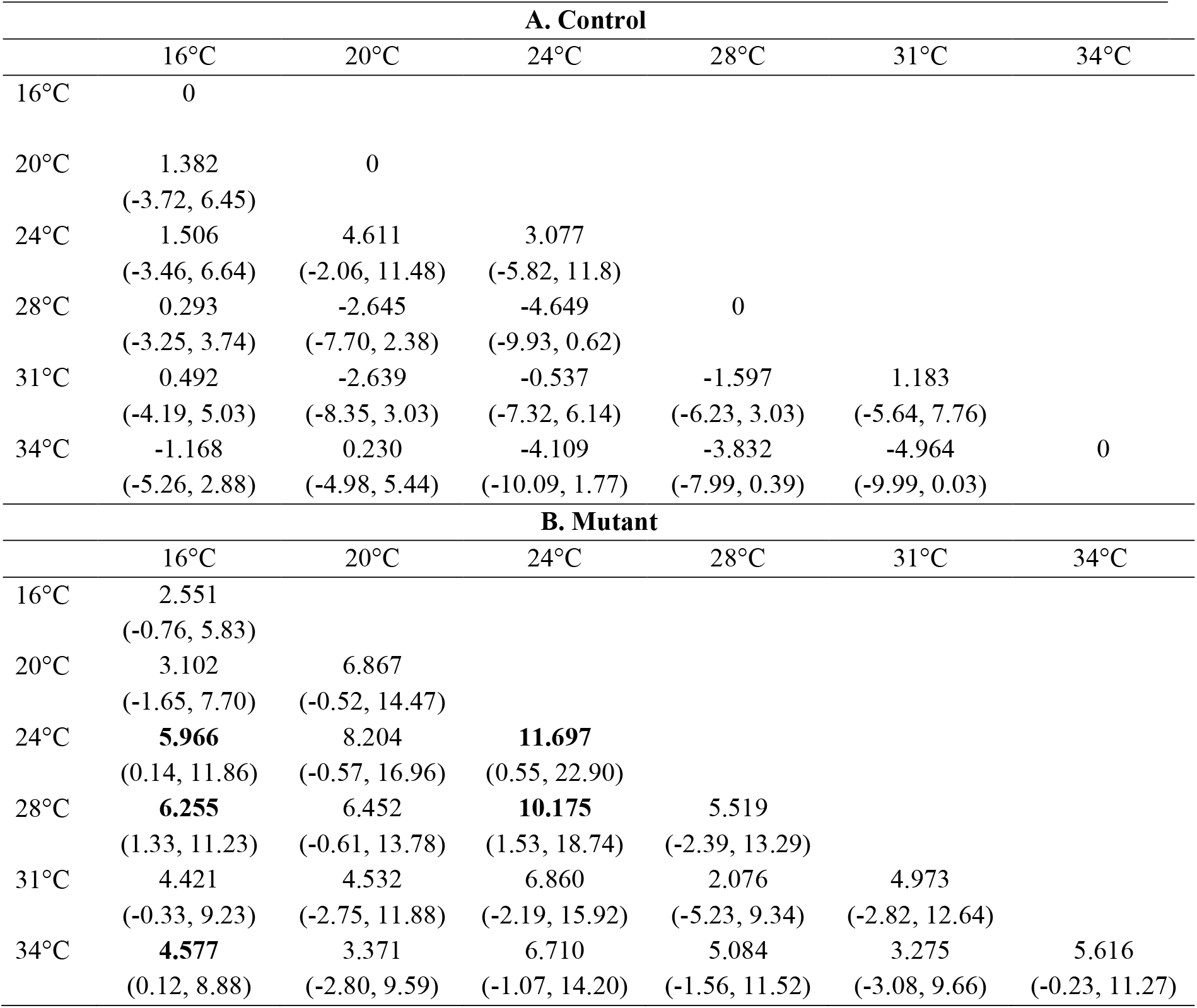
The among-family variance-covariance matrix for Control (A) and Mutant (B) treatments. The REML-MVN confidence intervals are reported below their respective estimates (90% CI for the variances on the diagonal, a one-tailed test against zero; 95% for covariances on the lower off-diagonal, a two-tailed test against zero). Parameters whose CI do not include zero are shown in bold. Estimates are from a model fitting an unconstrained among-family variance-covariance matrix. No CI could be estimated where the REML estimate was zero.

| **A. Control** |  |  |  |  |  |  |
| --- | --- | --- | --- | --- | --- | --- |
|  | 16°C | 20°C | 24°C | 28°C | 31°C | 34°C |
| 16°C | 0 |  |  |  |  |  |
| 20°C | 1.382<br>(-3.72, 6.45) | 0 |  |  |  |  |
| 24°C | 1.506<br>(-3.46, 6.64) | 4.611<br>(-2.06, 11.48) | 3.077<br>(-5.82, 11.8) |  |  |  |
| 28°C | 0.293<br>(-3.25, 3.74) | -2.645<br>(-7.70, 2.38) | -4.649<br>(-9.93, 0.62) | 0 |  |  |
| 31°C | 0.492<br>(-4.19, 5.03) | -2.639<br>(-8.35, 3.03) | -0.537<br>(-7.32, 6.14) | -1.597<br>(-6.23, 3.03) | 1.183<br>(-5.64, 7.76) |  |
| 34°C | -1.168<br>(-5.26, 2.88) | 0.230<br>(-4.98, 5.44) | -4.109<br>(-10.09, 1.77) | -3.832<br>(-7.99, 0.39) | -4.964<br>(-9.99, 0.03) | 0 |
| **B. Mutant** |  |  |  |  |  |  |
|  | 16°C | 20°C | 24°C | 28°C | 31°C | 34°C |
| 16°C | 2.551<br>(-0.76, 5.83) |  |  |  |  |  |
| 20°C | 3.102<br>(-1.65, 7.70) | 6.867<br>(-0.52, 14.47) |  |  |  |  |
| 24°C | **5.966**<br>(0.14, 11.86) | 8.204<br>(-0.57, 16.96) | **11.697**<br>(0.55, 22.90) |  |  |  |
| 28°C | **6.255**<br>(1.33, 11.23) | 6.452<br>(-0.61, 13.78) | **10.175**<br>(1.53, 18.74) | 5.519<br>(-2.39, 13.29) |  |  |
| 31°C | 4.421<br>(-0.33, 9.23) | 4.532<br>(-2.75, 11.88) | 6.860<br>(-2.19, 15.92) | 2.076<br>(-5.23, 9.34) | 4.973<br>(-2.82, 12.64) |  |
| 34°C | **4.577**<br>(0.12, 8.88) | 3.371<br>(-2.80, 9.59) | 6.710<br>(-1.07, 14.20) | 5.084<br>(-1.56, 11.52) | 3.275<br>(-3.08, 9.66) | 5.616<br>(-0.23, 11.27) |

We compared the magnitude of among-family variance following mutagenesis (Mutant minus Control) to published estimates from spontaneous mutation accumulation studies on traits classified by Conradsen et al. (2022) as physiological; these traits have an average magnitude of variance intermediate between fitness (life-history) and morphological traits (Figure 4b of Conradsen et al. 2022). We expanded the dataset to include estimates of *Drosophila serrata* locomotor activity at all temperatures assessed by Latimer et al. (2011). Mutational variance in *U*_*sprint*_ in the current study was within the range observed for these published estimates but was biased toward higher values (Figure 2).

**Figure 2.**
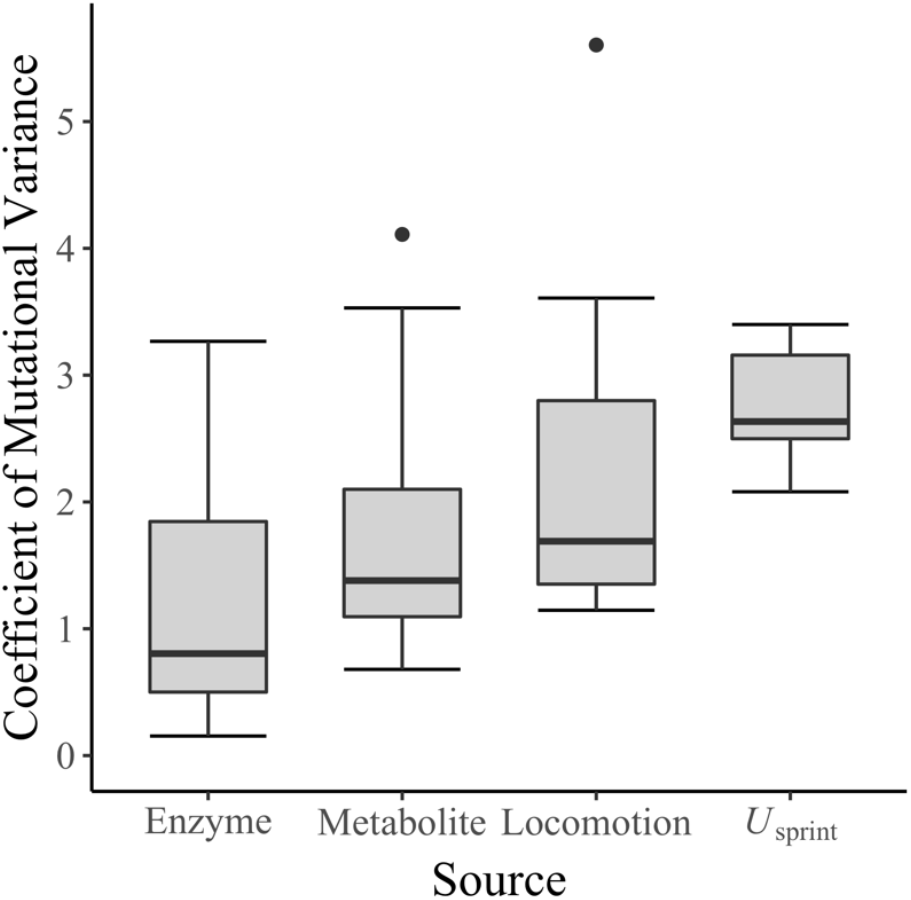
Comparison of the magnitude of mutational variance. Estimates of mutational variance were placed on a coefficient of variance scale: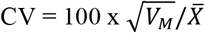where *V*_*M*_ was the mutational variance and 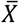 was the trait mean. Plotted are the median (black band), interquartile range (IQR; box) and three times the IQR (whiskers) plus any more extreme observations (circles). The 27 estimates of enzyme activity come from two studies in *Drosophila melanogaster* (Clark et al. 1995; Harada 1995), while the 27 estimates of metabolite pool size come from one study in *Caenorhabditis elegans* (Davies et al. 2016), derived as detailed Table S1 of Conradsen et al. (2022). The 16 locomotion estimates include male and female *D. serrata* activity measured at each of six temperatures (Table 1 of Latimer et al. 2014) as well as turn rate and velocity in *C. elegans* (Ajie et al. 2005; Estes et al. 2005) (derived as described in Table S1 of Conradsen et al. 2022). The six *U*_*sprint*_ estimates from the current study are the difference in among-family variance between Mutant and Control treatments, scaled by the Mutant treatment mean speed at that temperature.

The factor-analytic test of dimensionality best supported zero dimensions of among-family variance in the Control treatment (Table 2A). However, the REML-MVN CI of the eigenvalues supported two non-zero eigenvalues (Table 2A). Sztepanacz and Blows (2017) demonstrated REML-MVN CI are less robust to sampling error than factor analytic modelling and can lend statistical support to spurious covariance. While non-zero among-family variance in speed was estimated at two temperatures (24°C and 31°C: Table 1A), the model was unable to estimate (positive) variance at the other four temperatures and, further, the 90% CI of the among-family variance at both 24°C and 31°C spanned a wide range of negative through positive values (Table 1A) suggesting no statistical support for among-family variance in speed at any temperature. Reflecting this, the among-Control-family matrix was very ill-conditioned: the first eigenvalue (9.9; Table 2A) was substantially larger than the trace of the matrix (sum of the diagonal in Table 1A = 4.3), and there was a negative eigenvalue of nearly the same magnitude as the first (positive) eigenvalue (−9.3; Table 2). Therefore, we conclude that the weight of evidence is consistent with no statistical support for genetic variation in swimming speed (*U*_*sprint*_) in this WIK population.

**Table 2.** Dimensionality of among-family covariance matrices for A) Control and B) Mutant treatments. The results from fitting a nested series of reduced rank co-variance matrices at the among-family level are reported, where the Factor number corresponds to the number of dimensions modelled. Differences in the log-likelihood ratio between sequential models follows a chi-square distribution, with the degrees of freedom (df) corresponding to the difference in the number of parameters between the models. The Akaike Information Criterion (AIC) is also reported (with the smallest AIC, indicating the best fit model, shown in bold). The eigenvalues of the unconstrained among-family matrix estimated in each treatment (Table 1) are also reported, along with the REML-MVN 90% CI. In the Control treatment, the model constrained to six dimensions did not properly converge.

| Factor | # | -2log- | AIC | $\chi^2$ | df | P-value | Eigenvector | Eigenvalue | 90% CI |
| --- | --- | --- | --- | --- | --- | --- | --- | --- | --- |
|  | Parameters | likelihood |  |  |  |  |  |  |  |
| **A. Control** |  |  |  |  |  |  |  |  |  |
| 0 | 42 | 12043.4 | **12121.4** |  |  |  |  |  |  |
| 1 | 48 | 12031.9 | 12121.9 | 11.5 | 6 | 0.0740 | 1 | 9.884 | (3.54, 16.33) |
| 2 | 53 | 12021.5 | 12121.5 | 10.4 | 5 | 0.0647 | 2 | 6.996 | (2.81, 11.16) |
| 3 | 57 | 12021.3 | 12127.3 | 0.1 | 4 | 0.9979 | 3 | 1.655 | (-2.07, 5.37) |
| 4 | 60 | 12021.3 | 12133.3 | <0.0 | 3 | 0.9995 | 4 | -0.770 | (-3.72, 2.09) |
| 5 | 62 | 12021.3 | 12135.3 | <0.0 | 2 | >0.9999 | 5 | -4.175 | (-8.78, 0.54) |
| 6 | 63 | NA | NA | NA | NA | NA | 6 | -9.331 | (-18.85, -0.18) |
| **B. Mutant** |  |  |  |  |  |  |  |  |  |
| 0 | 42 | 11913.6 | 11997.6 |  |  |  |  |  |  |
| 1 | 48 | 11900.1 | **11992.1** | 13.5 | 6 | 0.0364 | 1 | 35.201 | (11.57, 58.6) |
| 2 | 53 | 11898.2 | 12000.2 | 2.0 | 5 | 0.8550 | 2 | 3.462 | (-1.20, 8.14) |
| 3 | 57 | 11896.8 | 12004.8 | 1.4 | 4 | 0.8475 | 3 | 3.173 | (-0.69, 7.03) |
| 4 | 60 | 11896.8 | 12012.8 | <0.0 | 3 | 0.9993 | 4 | 0.361 | (-3.29, 4.06) |
| 5 | 62 | 11896.8 | 12016.8 | <0.0 | 2 | >0.9999 | 5 | -0.502 | (-4.94, 4.00) |
| 6 | 63 | 11896.8 | 12014.8 | <0.0 | 1 | 0.9261 | 6 | -4.474 | (-9.52, 0.46) |

In the Mutant treatment, there was statistical support for one-dimension of among-family variance (Table 2B), suggesting either that mutation had introduced variance in *U*_*sprint*_ at only one assayed temperature (consistent with GxE), or that the effects of mutations were concordant across all assayed temperatures (consistent with no GxE). Although among-family variance was only statistically distinct from zero at 24°C, 90% CI estimates at three other temperatures were strongly skewed toward positive values, while the 90% CI of variance estimates substantially overlapped across the six temperatures (Table 1B). All pairwise covariances were positive (although only 26% were statistically distinct from zero: Table 1B). The first eigenvalue (35.2; Table 2B) was three times larger than the largest variance at any individual temperature and accounted for 95% of the total among-family variance. The contribution of speed at each temperature to this axis of variation (i.e., the eigenvector loadings: Table 3) were all in the same direction and of similar magnitude. Therefore, we suggest the data provide evidence that mutations have consistent effects on speed irrespective of the assay temperature, with no evidence of environment-specific effects.

**Table 3.** Major axes of among-family variance in swimming speed. The eigenvector loadings for the first and second axes of the unconstrained estimate of among-family variance in the Control (i.e., *g*_*max*_ and *g*_*2*_) and Mutant (i.e., *m*_*max*_ and *m*_*2*_) treatments are shown. The corresponding eigenvalues are reported in Table 2.

|  | Control |  | Mutant |  |
| --- | --- | --- | --- | --- |
| | $g_{max}$ | $g_2$ | $m_{max}$ | $m_2$ |
| 16°C | 0.184 | -0.149 | 0.317 | 0.252 |
| 20°C | 0.482 | 0.232 | 0.394 | -0.537 |
| 24°C | 0.737 | -0.130 | 0.591 | -0.064 |
| 28°C | -0.410 | -0.239 | 0.434 | 0.274 |
| 31°C | -0.022 | -0.622 | 0.306 | -0.478 |
| 34°C | -0.147 | 0.681 | 0.336 | 0.584 |

## Discussion

We detected phenotypic effects of mutation on a complex trait, swimming speed, but did not detect any mutational variance for the plasticity of speed in response to heterogeneity in water temperature. Mutations had concordant effects across the 18°C thermal gradient over which speed itself varied by more than 35cm^-1^ second. Our results are consistent with mutation introducing variation along a vertical shift, or faster-slower mode of thermal performance variation (Huey and Kingsolver 1989; Izem and Kingsolver 2005). In the only other study that we are aware of investigating mutational variance in locomotor thermal performance, Latimer et al. (2014) characterised the contribution of spontaneous mutation to variance in locomotor activity of *Drosophila serrata*, and similarly reported that most (76% or 70% in males or females, respectively) mutational variance was associated with a faster-slower mode. This mode of variation is interpreted as reflecting pleiotropic mutations with consistent direction of effects across all temperatures (Kingsolver et al. 2001), where selection for increased (or decreased) performance at one temperature would lead to correlated evolution at all temperatures, and under thermally heterogeneous conditions, consistent selection would effectively fix (eliminate) variants.

In contrast to mutation predominantly influencing average performance (Latimer et al. 2014), <1% of standing genetic variation in *D. serrata* was associated with the faster-slower mode; rather, most variation was aligned with a specialist-generalist mode (where the width of the performance curve varies) (Latimer et al. 2011). Other studies of standing genetic variation of thermally dependence in non-locomotor traits have also suggested little genetic variation associated with the faster-slower mode (e.g., Izem and Kingsolver 2005). Notably, mutational correlations among life-history traits are typically more strongly positive than the corresponding standing genetic correlation, reflecting selective elimination of concordantly deleterious mutations (Houle et al. 1994; Estes and Phillips 2006; McGuigan et al. 2011). Latimer et al. (2014) similarly suggested that the mismatch of mutational and standing genetic variation might indicate that most mutations affecting performance are deleterious, and do not persist in standing genetic variation. However, other studies of standing genetic variation in thermal performance curves for individual or population-level growth have provided contradictory evidence, suggesting that most (rather than least) variation is associated with a faster-slower mode (Yamahira et al. 2007; Moghadam et al. 2020), and several recent studies of locomotor performance found statistical support for heritability only of curve height (i.e., faster-slower variation) and not optimal temperature or curve width (Logan et al. 2018; Martins et al. 2019). To resolve the contrary predictions that standing genetic variation reflects selective process (Latimer et al. 2014) versus mutational limits (Yamahira et al. 2007) further data on the phenotypic variation introduced by mutation to thermal performance traits will be required. The distribution of mutations may influence molecular (Cano et al. 2022) and phenotypic (Houle et al. 2017) adaptation, and understanding whether standing genetic variation reflects limited input of the type of variation that would support local versus global adaptation is necessary for predicting adaptive responses to changing thermal conditions.

We observed no evidence that mutations had a biased direction of effect on swimming speed, consistent with a previous investigation of the effects of ENU-induced mutation on WIK zebrafish prolonged swimming speed at 28°C (McGuigan and Aw 2017). For fitness, mutation is predicted to be, and empirically supported as, typically biased toward lower values (Keightley and Lynch 2003; Keightley and Eyre-Walker 2007; Halligan and Keightley 2009). However, theoretical models of the maintenance of genetic variance in non-fitness traits typically assume no overall directional trend in the effects of mutations (Barton 1990; Kondrashov and Turelli 1992; Johnson and Barton 2005; Martin and Lenormand 2006); deviation from this expectation could result in substantial directional selection on traits (Zhang and Hill 2008). ENU-induced mutations in guppies decreased the rate of courtship displays (Herdegen and Radwan 2015), while spontaneous mutations decreased *D. melanogaster* larval crawling speed, adult heat tolerance, overall coordination (Huey et al. 2003) and male escape speed (Shabalina et al. 1997), and decreased velocity of *Caenorhabditis elegans* (Ajie et al. 2005). In contrast, spontaneous mutations decreased *D. serrata* locomotor activity only at the hottest assayed temperatures, increasing activity across a wide range of cooler temperatures (Latimer et al. 2014). Other performance traits (e.g., feeding rate and adult walking speed: Huey et al. 2003) showed no shift in trait mean under mutation accumulation. Given that these studies of whole-organism performance traits have reported mutational effects that span the full spectrum, further empirical and theoretical consideration of the evolutionary genetic consequences of non-symmetrical mutations for non-fitness traits is required. Differences among traits and studies may simply reflect unpredictable stochastic effects of sampling from the same, complex, distribution of effects. However, several factors, including study design, the traits themselves, genetic background, distribution of dominance, and the environmental range considered, may influence the distribution of phenotypes generated by mutation.

A challenge faced by classical multi-generational mutation accumulation experiments is evolution of the ancestor (discussed in Lynch et al. 1999). Some heterogeneity in direction of effect may reflect estimation errors arising from changes in the ancestor. This problem is avoided using mutagenesis, where the unmutated comparison population has not had time to evolve. Notably, while classical mutation accumulation experiments are typically initiated from a single ancestral genotype (inbred line), mutagenesis experiments introduce mutation to multiple (here, 50) genetic backgrounds simultaneously. Spontaneous mutation rate (Sharp and Agrawal 2012; Schrider et al. 2013; Huang et al. 2016) and distribution of mutational fitness effects (Fisher 1930; Orr 1998; Silander et al. 2007; Stearns and Fenster 2016) are influenced by ancestral genotype. Whether bias in the direction of mutational effects on non-fitness traits can be (partially) explained by genetic background, or the contribution of the trait to fitness, remains to be determined.

Observation of both thermally dependent and directionally biased effects of mutations may also be influenced by dominance. We focused here on heterozygous effects: new mutations will be expressed only in heterozygotes due to their rarity, and it is therefore their heterozygous effect that will determine the selection they experience. Our observation that mutation significantly increased genetic (among-family) variance in speed indicates that at least some induced mutations were not fully recessive in their effects on the focal trait. Mutations with larger homozygous deleterious effects on fitness tend to be more recessive (Agrawal and Whitlock 2011), but the joint distribution of dominance coefficients of pleiotropic mutations affecting fitness and other traits is not well characterised. Mutations causing notable defects in larval swimming performance have been reported to be typically homozygous lethal (Granato et al. 1996), suggesting directional dominance on fitness is correlated with directionality of effects on speed for these large-effect mutations. Similarly, a study of a large cohort of racehorses found evidence of a negative correlation between an individual’s inbreeding coefficient and race performance, consistent with correlation of directional dominance of fitness with effects on performance (Todd et al. 2018). Correlated environment-dependent reversals in dominance (recessive to dominant) and fitness (deleterious to beneficial) effects of mutations are predicted to accelerate adaptation (Muralidhar and Veller 2022). Here, the observed concordance of mutational effects on speed are not consistent with environment-dependent changes in the dominance coefficient. Nonetheless, our data cannot exclude the possibility of an unobserved class of undetected mutations with recessive effects that had directionally biased, or non-concordant pleiotropic effects (antagonistic or conditionally neutral), shifting mean speed and generating genotype-by-environment interaction variation.

We assayed a set range of temperatures (16°C – 34°C), and while speed varied markedly with temperature, it was nonetheless notable that average speed changed very little between 28°C and 34°C (Figure 1). We cannot exclude the possibility that the effects of the sampled mutations on swimming speed were different outside the considered thermal range. The frequency distribution of fitness effects may be shifted under hotter temperatures (Xu 2004; Chu et al. 2020). However, direct effect of temperature on population growth rate in microbes, where population size limits (bottlenecks) are not imposed per generation, may allow greater opportunity for selection to shift the observed frequency of mutations at hot relative to cool temperatures (Wahl and Agashe 2022), with stronger selection on deleterious mutations under hotter conditions (Berger et al. 2021).

Consistent with Latimer et al. (2014), we assessed adult phenotypes under acute exposure to different temperatures. The timing and duration of exposure to heterogeneous environments can affect thermal performances (Rezende et al. 2014; Kellermann et al. 2019; Pottier et al. 2022), and the observed distribution of mutational effects may likewise change. Fully replicating multiple rearing temperatures to impose long-term exposure, (e.g., through larval development) is logistically limiting. Further, temperature-dependent effects on larval viability may result in the phenotypic effects of different mutations being sampled at different temperatures. Selection acting among siblings within a family of an ENU mutated sire can affect the frequency of mutations within families and observation of their phenotypic effects (Walsh and McGuigan 2018). Temperature also affects development rate in ectotherms, potentially result in confounding of the effects of mutation, development and temperature. We previously showed swimming speed in adult zebrafish changes with age, independent of size or known maturation and senescence boundaries (McGuigan and Aw 2017). While natural populations encounter varying temperatures throughout life, disentangling how genetic effects cause plastic responses to that variation is challenging.

Our results by no means suggest the absence of any class of mutations with non-concordant pleiotropic effects across temperatures. Quantitative genetic parameters are notoriously difficult to accurately estimate, given their relatively large sampling error (Klein et al. 1973; Klein 1974). Notably, standing genetic variance for thermal performance traits was below statistical detection limits in several recent studies (Driessen et al. 2007; Logan et al. 2018; Martins et al. 2019; Logan et al. 2020; Bodensteiner et al. 2021). Our results (limited statistical support for heritable variation per temperature, but strong support for heritable variation in speed when considering all data) suggest that, given the pervasive presence of temperature-independent mutational effects, we benefited from repeated measures of the same phenotype to improve estimation. A recent investigation suggested that sampling error is likely to make a non-negligible contribution to heterogeneity among estimates of mutational variance (Conradsen et al. 2022). Similarly, a recent meta-analysis of population means of thermal physiological limit traits suggested that < 8% of 428 estimates were supported by sufficient sample sizes for the mean to be estimated with a high-level of accuracy (Duffy et al. 2021). Thermally dependent traits are likely to remain the focus of understanding how populations can persist under and adapt to changing conditions, but we suggest that the experimental effort involved in obtaining useful and robust parameter estimates is far from trivial, while the risk of inaccurate estimation must also be carefully considered. In the current study, we collected a total of 3,497 measures of speed, which took 677 person hours to record (without accounting for time taken for acclimating fish prior to trials, or any aspect of breeding and husbandry). Without additional information on the relative frequency and effect size of mutations with thermally dependent effects it is not simple to predict how much larger an experiment will be required to detect, above measurement error, and characterise other, non-concordant patterns of variation across temperatures. The apparent benefit of repeated measures suggests that improved precision of measurement of performance (via within individual and within family replication) will be important, along with increasing the number of genotypes (families), which would support sampling of more mutations.

Mutagenesis could greatly expand the range of taxa in which the distribution of mutational effects on phenotype (and fitness) can be explored. The mutational variance we observed for sprint speed was greater than the per-generation rate of mutational input reported from classical mutation accumulation (MA) studies of related traits (Figure 2), suggesting that we had induced more mutations than typically sampled in long-running MA experiments (although differences in effect size or the greater mutational target size of speed than an individual enzyme or metabolite may also have contributed). Ideally, as is possible in microbial systems, phenotypic effects of individual mutations would be considered. However, isolating individual mutations to characterise their effect remains challenging in higher eukaryotes. Genome sequencing of multicellular eukaryote MA lines, maintained over tens to hundreds of generations, typically reveal multiple (∼20 – 90) mutations per line accumulating over these time frames (Schrider et al. 2013; Huang et al. 2016; Assaf et al. 2017; Flynn et al. 2017). When MA lines (or here, families) diverge from one another at multiple loci, strong correlations among traits (here, speed at the different temperatures) can arise through both heterogeneity of mutation number among lines or via linkage, but such spurious correlations depend on mutations having biased, unidirectional, effects on the traits (Keightley et al. 2000). However, co-segregation of mutations having opposing effects on the same traits may have obscured patterns arising from more complex (antagonistic or conditionally independent) pleiotropic effects. More generally, epistatic interactions among mutations may influence the observed patterns of phenotypic variation, as apparent in the effect of genetic background on the frequency of beneficial mutations (Silander et al. 2007; Perfeito et al. 2014; Stearns and Fenster 2016). In the current experiment, there was potential for epistatic interactions to influence observed patterns of phenotypic variation both due to among-sire differences in their genotype (i.e., genetic background), and due to the specific set of mutations induced in each sire.

Further information is also required on how well the induced mutations reflect the spectrum of naturally arising mutations (Katju and Bergthorsson 2019). The zebrafish genome is large (1400 Mb) relative to other multicellular eukaryotic taxa for which extensive mutational data exist (100 – 200 Mb for *D. melanogaster, Daphia pulex, Caenorhabditis* species and *Arabidopsis thaliana*), and, in contrast to these other models in which a genetically homogeneous ancestor can be rapidly established, the longer generation time of zebrafish (and other vertebrates) makes this more challenging. Novel mutations must therefore be accurately detected against a background segregating genetic variation (Coe et al. 2009; Brown et al. 2012; Butler et al. 2015; Suurvali et al. 2020). Thus, while studies of zebrafish (and other vertebrate models) can exploit considerable genomic tools, dissecting contributions from overall mutation number versus effect sizes will be challenging.

Heterogeneity in allelic effects among environmental contexts contribute to phenotypic diversity and adaptive potential of natural populations. The role of mutation in shaping the distribution of this diversity has the potential to influence direction and rate of evolution of populations. The limited amount of evidence to date suggests that mutations with concordant effects across thermal gradients are more frequent or of larger effect than mutations with antagonistic or conditionally neutral effects. However, the patterns of intra and inter-specific genetic diversity of thermally dependent traits suggest major contributions from genetic variants with non-concordant (antagonistic or conditionally neutral) effects. Further investigation of mutational effects is required to reconcile these observations. Studies addressing this knowledge gap will need to be large, ensuring that effects can be accurately determined. Mutagenesis may facilitate studies across more taxa, and reduce the logistical burden of long-term maintenance.

## Acknowledgments

We thank Nick Appleton, Ash Kannan, and Laura Davis for their contributions to data collection and fish husbandry. We also thank The University of Queensland Biological Resources Aquatics Team for their assistance with husbandry and maintenance of life-support systems. This work was funded by the Australian Research Council DP180101801 awarded to KM. The authors declare no conflicts of interest.

